# Intracellular cyclic AMP levels modulate differential adaptive responses on epimastigotes and cell culture trypomastigotes of *Trypanosoma cruzi*

**DOI:** 10.1101/677112

**Authors:** Tamara Sternlieb, Alejandra C. Schoijet, Guillermo D. Alonso

**Author notes:** Correspondence G. D. Alonso, Laboratorio de señalización y mecanismos adaptativos en tripanosomátidos, Instituto de Investigaciones en Ingeniería Genética y Biología Molecular “Dr. Héctor N. Torres”, Vuelta de Obligado 2490 (C1428ADN), Ciudad Autónoma de Buenos Aires, Argentina.

## Abstract

Among the many environmental challenges the parasite *Trypanosoma cruzi* has to overcome to complete its life cycle through different hosts, oxidative stress plays a central role. Different stages of this parasite encounter distinct sources of oxidative stress, such as the oxidative burst of the immune system, or the Heme released from hemoglobin degradation in the triatomine’s midgut. Also, the redox status of the surroundings functions as a signal to the parasite, triggering processes coupled to differentiation or proliferation. Intracellular second messengers, like cAMP, are responsible for the transduction of environmental queues and initiating cellular processes accordingly. In trypanosomatids cAMP is involved in a variety of processes, including proliferation, differentiation, osmoregulation and quorum sensing. Trypanosomatid phosphodiesterases (PDE) show atypical pharmacological properties and some have been involved in key processes for the survival of the parasites, which validates them as attractive therapeutic targets. Our work here shows that cAMP modulates different processes according to parasite stage. Epimastigotes become more resistant to oxidative stress when pre-treated with cAMP analogs, while trypomastigotes do not alter their response to oxidative stress under the same treatment. However, cAMP analogs do increase trypomastigotes infectivity *in vitro*. Also, we show that TcrPDEA1, a functionally enigmatic phosphodiesterase with very high Km, is involved in the epimastigotes response to oxidative stress.

## Introduction

*Trypanosoma cruzi* is the causative agent of Chagas disease. As a parasitic unicellular organism, it has developed specific mechanisms to allow it to cope with the extreme and sudden changes in the environment as it is transmitted from an insect vector to the mammalian host and vice versa. A testimony of this adaptability is the presence of at least three different developmental stages during its life cycle, some replicative but not infective (amastigote and epimastigote) and other infective but not replicative (metacyclic and bloodstream trypomastigote). Despite the high relevance of this rapid adaptation to environmental changes to the progression of the trypanosomatid life cycles, little is known about the signals and molecules that trigger the stage specific metabolisms and differentiation from one stage to the other.

Cyclic AMP is a vital second messenger in several transduction pathways in eukaryotic cells. In trypanosomatids it has been implicated in stage differentiation, proliferation, stress response and even social motility since three decades ago (Bao et al., 2008; Bhattacharya et al., 2008; Biswas et al., 2011; D’Angelo et al., 2004; Fraidenraich et al., 1993; Gonzales-Perdomo et al., 1988; Laxman and Beavo, 2007; Makin and Gluenz, 2015; Rangel-Aldao et al., 1988; Tagoe et al., 2015). Initially, higher intracellular cAMP and *T. cruzi* Adenylyl cyclase (AC) activity were shown to stimulate metacyclogenesis (Fraidenraich et al., 1993; Gonzales-Perdomo et al., 1988; Rangel-Aldao et al., 1988). Additional studies demonstrated that elevated intracellular cAMP could arrest DNA, RNA and protein synthesis (Santos and Oliveira, 1988), which concomitantly led to a proliferation arrest in epimastigotes. Later, a new family of phosphodiesterases (PDEs) was identified in trypanosomatids, and they were connected with certain stress response transduction pathways (Alonso et al., 2007; Bhattacharya et al., 2009), and in *T. brucei* TbPDEB1 and TbPDEB2 have even been validated as drug targets (Bland et al., 2011; De Koning et al., 2012; Oberholzer et al., 2007; Torphy and Page, 2000). Interestingly, very few cAMP effector proteins, besides protein kinase A (PKA) have been identified (Jäger et al., 2014), and the proteins linking the activation of AC or PKA and the consequences observed in phenotype have not yet been uncovered. The relevance of this second messenger in *T. cruzi* increases since no orthologues for enzymes of the cGMP pathway are found in its genome (Naula and Seebeck, 2000).

In *T. cruzi*, some enzymes of the cAMP pathway have been annotated after the genomic sequencing of this parasite. An Adenylyl cyclase with a mammal Guanylyl cyclase like structure was identified and associated with starvation induced metacyclogenesis (Hamedi et al., 2015). From its structure (a C-terminal conserved intracellular catalytic domain, a single transmembrane helix, and a variable N-terminal domain of unknown function) it was hypothesized that it functions as an extracellular sensor, activating intracellular mechanisms in response to external stimulus (D’Angelo et al., 2002; Taylor et al., 1999; Torruella et al., 1986). It was shown to be involved in metacyclogenesis in *T. cruzi* and other processes like differentiation, quorum sensing and social motility in *T. brucei* (Hamedi et al., 2015; Lopez et al., 2015; Oberholzer et al., 2015, 2010). Several PDEs have been identified in trypanosomatids, all belonging to Class I, and some have been validated as drug targets (Alonso et al., 2007; Oberholzer et al., 2007). PDEA, of *L. donovani*, is a cytosolic enzyme differentially regulated through the parasite’s life cycle and involved in the oxidative stress response (Bhattacharya et al., 2009). PDEB1 and PDEB2 are membrane associated phosphodiesterases of *T. brucei* and are essential for its proliferation (Oberholzer et al., 2007). PDEC is involved in osmotic stress response in *T. cruzi*, specifically in the contractile vacuole complex (Schoijet et al., 2011). Protein kinase A regulatory and catalytic subunits are also present in *T. cruzi* with a cAMP specific dependent kinase activity (Bao et al., 2008; Huang et al., 2006, 2002). However, in spite of novel proteins with the ability to bind cyclic nucleotides being identified in *T. cruzi* (Jäger et al., 2014), how cAMP and the related enzymes act on all these different processes is still to be explained.

Oxidative stress plays a major role in several stages during *T. cruzi* life cycle. After the epimastigotes enter the vector’s midgut, they encounter large amounts of heme released from hemoglobin degradation and other oxidative products of blood digestion (Machado-Silva et al., 2016). As the digestive tract approaches to the urinary tract of the insect, the environment changes its redox status to a more reductive than oxidative one. Nogueira et. al. proposed that the redox status of the environment acts as one of the signals that determine parasite proliferation or differentiation (Nogueira et al., 2015). They proved that reductive species are capable of arresting epimastigote proliferation and stimulate metacyclogenesis, while proliferation requires an oxidative environment, *in vitro* and *in vivo* (in the insect host). Furthermore, they had already showed that heme-induced ROS was responsible for the proliferation induction and H_2_O_2_ was the signaling molecule, while also suggesting a mechanisms involving CaMKII (de Almeida Nogueira et al., 2011). The insect vector also holds an immune response that is activated in response to trypanosomatid infections, and it has been shown that nitric oxide (NO) and reactive oxygen intermediates are an important part of it (Ursic-Bedoya and Lowenberger, 2007).

Upon entering the mammal host, the parasite must cope with and evade the immune attack. One of the innate immune early responses is the oxidative burst, where reactive oxygen and nitrogen species (ROS and RNS) generate oxidative stress on the trypomastigote. Oxidative stress appears to be necessary for an optimal infectivity. Paiva et al showed that ROS contribute to amastigote growth inside macrophages and increase overall parasitism (Paiva et al., 2012). Later on, Goes et al also showed that H_2_O_2_ in low concentrations acts as a specific signal for the trypomastigotes that promotes greater proliferation in macrophages (Goes et al., 2016). The identification of the parasite mechanisms of evasion of the immune system and the proteins involved could reveal new drug targets for chemotherapy. This is of utter importance, since current chemotherapies present toxicity for the patients and are ineffective to cure the disease once the chronic phase is established. Moreover, there has been increased resistance of the parasites to the currently used drugs (Alsford et al., 2013).

Bhattacharya et al have demonstrated that in *Leishmania donovani* the cytosolic phosphodiesterase LdPDEA has a central role in the oxidative stress response and infectivity (Bhattacharya et al., 2009). Incubation with cAMP analogs or knock-down of the PDEA both elicited promastigotes to be more resistant to ROS and RNS *in vitro*. Overexpression of PDEA, on the other hand, made parasites less infective in a macrophage culture. These and other results led them to theorize that regulation of expression level of the PDEA during *L. donovani* transition from the insect vector to the vertebrate host allowed the parasite to make a shift in the utilization of the intracellular antioxidant species pool.

In this paper, we show evidence for the first time that the cAMP transduction pathway is involved in the oxidative stress response in *T. cruzi* epimastigotes. We demonstrated that cAMP analogs confer protection to epimastigotes under oxidative stress conditions, allowing survival and proliferation. We also investigate which PDEs are involved in this resistance process using PDE overexpressing parasites. Finally, we evaluate the effect of oxidative stress in combination with a cAMP analog on trypomastigotes through short infections on Vero cells.

## Materials and Methods

### Chemicals and reagents

All radiochemicals used in this work were purchased from Perkin Elmer Life Sciences, and restriction endonucleases were from New England Biolabs, Beverly, MA. Bactotryptose, yeast nitrogen base, and liver infusion were from Difco. All other reagents were purchased from Sigma.

### Bacterial, Mammal and Parasite Cell Lines and Cultures

*E. coli* bacteria of the strain DH5αF’ (F’/endA1 hdR17 (rk-mk+) supE44 thi-1 recA1 gyrA96 (Nalr) relA1 Δ(lacZYA-argF)U169 (m80lacZΔM15)) were used for replication of plasmid constructs. *T. cruzi* epimastigotes of the CL Brener strain were cultured at 28 °C for 7 days in liver infusion tryptose (LIT) medium (5 g l^−1^ liver infusion, 5 g l^−1^ bacto-tryptose, 68 mM NaCl, 5.3 mM KCl, 22 mM Na2HPO4, 0.2% (W/V) glucose, and 0.002% (W/V) hemin) supplemented with 10% (V/V) FCS, 100 U ml^−1^ penicillin and 100 mg l^−1^ streptomycin. Cell viability was assessed by direct microscopic examination. Trypomastigotes Tulahuen LacZ clone C4 (ATCC® PRA330(™)) were grown on monolayers of *Cercopithecus aethiops* (green monkey) Vero cells (ATCC CCL-81). Vero cells were grown in MEM medium containing 10% of SFB.

### Western blots

For Western blot analysis, proteins were solved in 10% (w/v) SDS-polyacrylamide gel electrophoresis as described by Laemmli (Laemmli, 1970) and electrotransferred to Hybond-C membranes (Amersham Pharmacia Biotech, Piscataway, USA). The membranes were blocked with 5% (w/v) non-fat milk suspension in TBS-Tween for at least 3 h. Blocked membranes were then incubated over night with a 1:1000 dilution of anti TcrPDEA1 polyclonal antibody (Alonso et al., 2007). Detection was carried out by incubating with a 1:5000 dilution of a goat anti-rabbit conjugated to peroxidase (Perkin Elmer and Sigma, Aldrich respectively). The latter were developed with the ECL Plus^™^ Western blotting detection system (PerkinElmer Life Sciences).

### Cyclic AMP phosphodiesterase assays

TcrPDEC2 activity was determined as described by Thompson and Appleman (Thompson et al., 1974) with the modifications introduced by Londesborough (Londesborough, 1977). The reactions were performed in the presence of 40 mM Tris–HCl pH 7.5, 10 mM Mg^2+^ and 0.5 μCi (20 nM) of [^3^H] cAMP. Incubations were carried out at 30 °C for 10 min in a total volume of 100 μl.

### Plasmid Constructs, Parasite Transfection and Transgenic Selection

We had previously obtained bacteria clones which replicate plasmid constructs in the pTREX vector containing the genes for three of the known *T. cruzi* PDEs: TcPDEA1 and TcPDEC2 (Alonso et al., 2007; D’Angelo et al., 2004; Schoijet et al., 2011). These plasmids where purified in large quantities using the Promega MIDI-prep kit and protocol and then transfected into three different CL Brener epimastigotes cultures, including a transfection control culture and a culture transfected with the pTREX empty vector, as described previously (Vazquez and Levin, 1999). Transfected lines were selected and maintained with G418 antibiotic (Gibco BLR, Carlsbad, CA) to a final concentration of 500 μg ml^−1^. After 60 days of treatment with 500 μg ml^−1^ of G418 stable cell lines were confirmed by enzymatic activity assay and Western blot.

### Oxidative Stress and Tritiated Thymidine Incorporation Assays

*T. cruzi* epimastigotes were grown to a concentration between 1 and 2 × 10^7^ parasites ml^−1^ (mid log phase), confirmed by microscopic cell counting. Culture was washed with PBS and resuspended in PBS supplemented with 2% (W/V) glucose. Hydrogen peroxide was applied in different concentrations to produce oxidative stress, and only PBS was added to a control culture. After a 10-minute incubation at 28 °C, cells where washed and resuspended in LIT medium containing 0.5 μCi ml^−1^ of tritiated Thymidine. Each culture was distributed in triplicates on a 96 well plate. Absorbance at 600 nm was measured at time 0 h and Thymidine incorporation was measured at 24 h after the assay. Proliferation percentages where calculated as following:

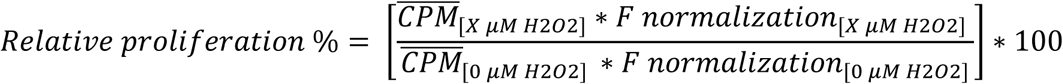

where ‘CPM’ is the media counts of each culture assayed under stress conditions (X μM) after 24 h, and the ‘0 μM’ represents the control culture which wasn’t exposed to hydrogen peroxide. The ‘F normalization’ factor is calculated with the absorbance at time 0 h to normalize the cultures to the number of parasites at the beginning of the assay. It is calculated as follows:

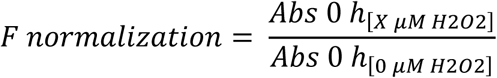

### Preincubations with cAMP Analogs

Permeable and stable cAMP analogs 8-bromoadenosine 3’:5’-cyclicmonophosphate (8-Br-cAMP) and 8-(4-Chlorophenylthio) adenosine 3’,5’-cyclic monophosphate (8-pCPT-cAMP) were used as pretreatments previous to oxidative stress. Different concentrations of the analogs (ranging from 0 μM to 1 mM) were applied to *T. cruzi* epimastigotes in LIT culture medium at 28 °C for 30 minutes, after which cultures were washed and resuspended in PBS-glucose 2% (W/V) for hydrogen peroxide treatment.

### In vitro *Infection Assays*

Protocol was applied as in Costa L. et. al. and Jeffrey Neitz R. et. al. (Costa et al., 2016; Jeffrey Neitz et al., 2015). To establish the ideal multiplicity of infection (MOI) for a 24 h infection, a Z factor was calculated according to the following equation:

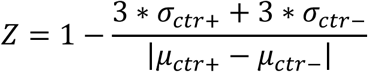

where μ is the mean and σ is the standard deviation of the positive control infection (ctr +) or the uninfected Vero cells, negative control (ctr -). MOIs 20:1, 10:1 and 5:1 were assayed and cell number of 1×10^4^ or 2×10^4^ per well, and the condition where the value of Z was between 0.5 and 1 was selected.

Vero cells were plated at 1×10^4^ cells per well on a 96 well plate so that three technical replicates of each infection were made. *T. cruzi* Tulahuen trypomastigotes overexpressing the *E. coli* β-galactosidase protein were treated with different combinations of cAMP analog and H_2_O_2_, resuspended in MEM culture medium supplemented with 3% SFB, and distributed in triplicates on the plated Vero cells. After 24 h incubation at 37 °C, culture medium containing trypomastigotes was extracted and replaced for fresh MEM-SFB 3% medium and the plate was incubated another 24 h at 37 °C. 48 h after infection started, cells and amastigotes were lysed with lysis buffer (25 mM Tris HCl pH 7.8, 2 mM EDTA, 2 mM DTT, 1% Triton x-100, 10% Glycerol) at 37 °C for 10 minutes, and reaction buffer 2x containing ortho-nitrophenyl-β-D-galactopyranoside (ONPG) (200 mM sodium phosphate buffer pH 7, 2mM MgCl_2_, 100 mM 2-Mercaptoethanol, 1.33 mg ml^−1^ ONPG) was added. Colorimetric changes were measured every hour by absorbance at 420 nm.

### Statistical Analysis

All statistical analysis were performed with GraphPad Prism 6.

## Results

### Dose dependent effect of hydrogen peroxide on epimastigote proliferation

The first environment that bloodstream trypomastigotes of *T. cruzi* reach after a triatomine’s blood meal is the anterior region of the midgut, and a few hours after, in the same region, they differentiate to epimastigotes. As epimastigotes pass through different sections of triatomines gut, this parasite form needs to proliferate and finally to differentiate to metacyclic trypomastigotes, when they reach the rectum of the vector.

In the anterior region of the midgut, large amounts of hemoglobin are digested producing an increase in the concentrations of heme, and the consequent increase in the formation of reactive oxygen species (ROS) (Ryter and Tyrrell, 2000). Considering that this is the initial environment that epimastigotes are obligated to deal with, and that the molecular mechanisms implicated in the oxidative stress response in *T. cruzi* are not fully understood, we decided to initiate our assays with epimastigotes. In order to reproduce similar conditions *in vitro*, we used hydrogen peroxide as an oxidative stress agent. To assess the stress caused by hydrogen peroxide on epimastigote cultures, and determine a dynamic range, we used an indirect proliferation measure method consisting of the incorporation of [H^3^]-Thymidine during DNA replication. We assayed a control condition with epimastigotes treated only with PBS and treatments with three different hydrogen peroxide concentrations. Treatments were applied for ten minutes at 28 °C. Afterwards hydrogen peroxide was washed and epimastigotes were resuspended in LIT culture medium containing [H^3^]-Thymidine. After 24 h, Thymidine incorporation was measured in a liquid scintillation counter. Results showed that for epimastigote cultures, 100 μM of hydrogen peroxide has no effect on proliferation, 150 μM has a sub-lethal effect (∼ 50 % proliferation), and 200 μM or higher concentrations impede recovery of proliferation after 48 h (Figure 1, and data not shown).

**Figure 1.**
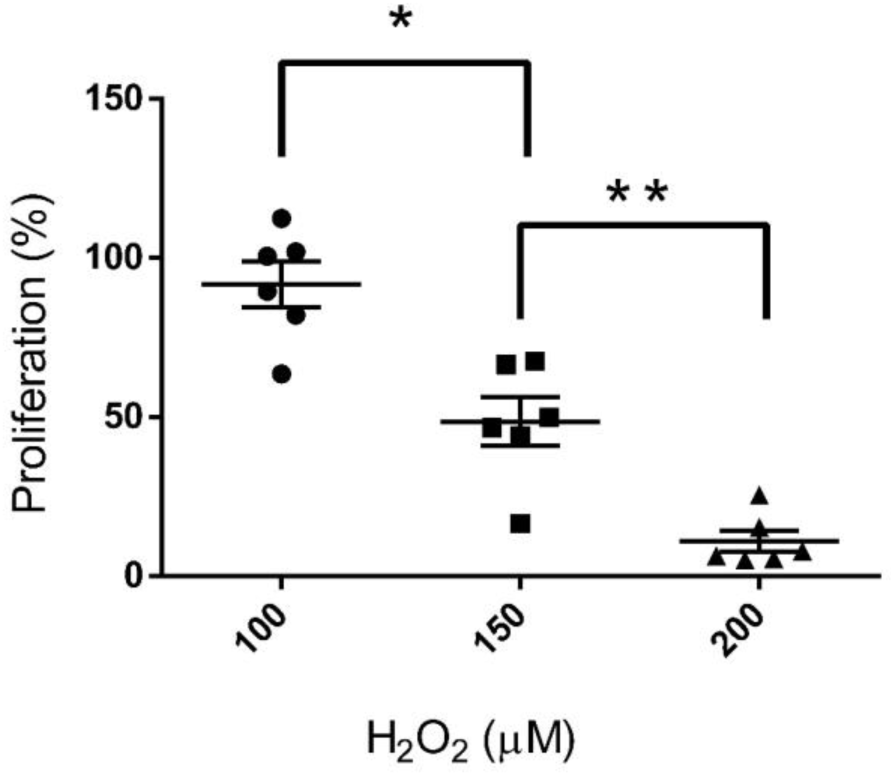
Dose dependent effect of hydrogen peroxide on epimastigote proliferation. Proliferation was measured as incorporation of [^3^H]-Thymidine in a 24 h culture time after treatment. Values are normalized to the control culture incubated only with PBS. One-way ANOVA with Tukey multiple comparison test indicated significant differences between effects (n = 6). (*) p<0.05 (**) p<0.01.

### Increase in intracellular cAMP enhances epimastigotes proliferation under oxidative stress conditions

It is very well documented that cAMP plays a crucial role in the differentiation of trypanosomatids (Fraidenraich et al., 1993; Gonzales-Perdomo et al., 1988; Rangel-Aldao et al., 1988) and in *Leismania donovani* it was reported that intracellular cAMP plays an important role in differentiation-coupled induction of oxidative damage resistance (Bhattacharya et al., 2008). Following that rational, we hypothesize that cAMP is also playing an important role in *T. cruzi* epimastigotes preventing the oxidative stress damage. To investigate whether an increase of intracellular cAMP could protect *T. cruzi* against oxidative stress, we assessed epimastigotes’ proliferation 24 h after they were treated alternatively with two different cAMP analogs (8Br-cAMP and 8-pCPT-cAMP) and, subsequently, stressed with 150 μM hydrogen peroxide. Our statistical analysis showed that an increase in cytosolic cAMP levels improved proliferation under oxidative stress conditions (p values 0.0714 and 0.0144, respectively). Depending on the analog concentration and its effect in the absence of stress, this improvement varied between ∽10% and ∽15% of Thymidine incorporation. Figure 2 shows that this effect is more significant when 8-pCPT-cAMP is applied. This could be due to 8-pCPT-cAMP higher lipophilicity, cellular permeability and stability. Furthermore, in non-stressed controls, pre-incubation with cAMP analog had no statistically significant effect on proliferation.

**Figure 2.**
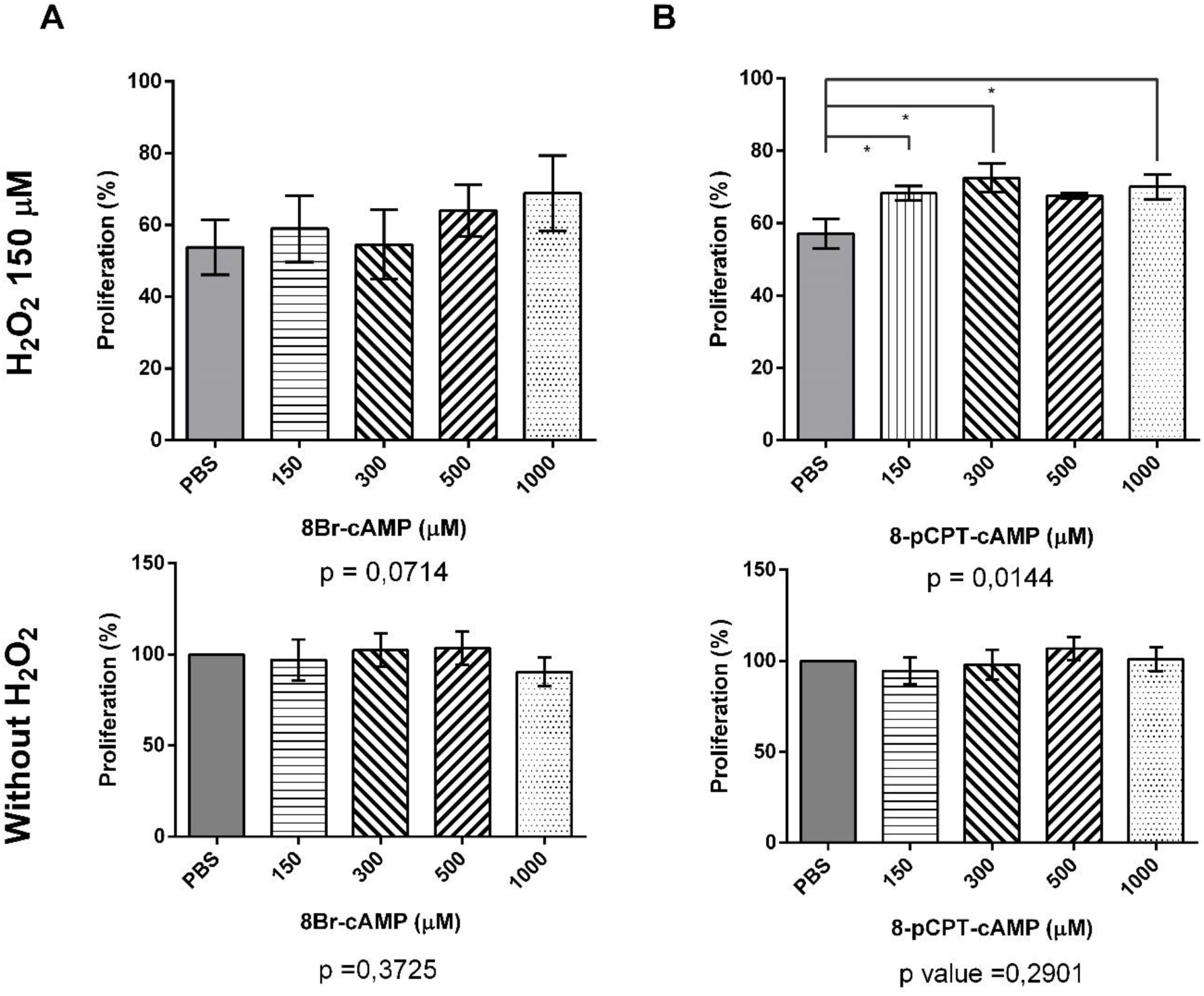
Stress response after pre-incubation with cAMP analogs. *T. cruzi* epimastigotes were incubated in LIT medium with different cAMP analog concentrations (8Br-cAMP for panel A and 8-pCPT-cAMP for panel B) for 30 minutes at 28 °C. cAMP analog was then removed, and parasites were incubated with 150 μM hydrogen peroxide in PBS-glucose 2% for 10 minutes at 28 °C (Upper panels). Hydrogen peroxide was removed and parasites were incubated for 24 h in LIT with [H^3^]-Thymidine. Finally, [^3^H]-Thymidine incorporation was measured and normalized to the culture treated with the corresponding cAMP analog concentration only (lower panels), obtaining a proliferation percentage. Statistical analysis consisted of a One-way ANOVA (n = 4), p values below graphs. Between treatments comparisons were analyzed by t-test. Error bars represent standard error of the mean. (*) p<0.05.

### Overexpression of TcrPDEA1, a cytosolic phosphodiesterase, decreases the resistance to oxidative stress in epimastigotes

Since pre-incubations with cAMP analogs showed to enhance epimastigotes proliferation under oxidative stress conditions, we decided to investigate whether the intracellular cAMP level, genetically modulated by the overexpression of TcrPDEA1, modifies the oxidative stress response capability of epimastigote forms. It is well documented that differentially distributed intracellular cAMP microdomains are regulated by cAMP specific phosphodiesterases with precise intracellular localizations (D’Angelo et al., 2004; Oberholzer et al., 2007; Schoijet et al., 2011). Additionally, in *Leishmania donovani* cAMP-PDE activities in the membrane and microsomal fractions remain constant during the parasite differentiation from log phase to the axenic amastigote stage, while in the cytosolic fraction the cAMP-PDE activity diminishes gradually (Bhattacharya et al., 2009). Moreover, in the same publication, Bhattacharya et al, suggest that the LdPDEA is the enzyme responsible for cytosolic cAMP reduction along the way of differentiation from promastigote to amastigote. To further evaluate this scheme in *T. cruzi*, we transfected epimastigotes with the pTREX vector containing the TcrPDEA1 (the gene ortholog to LdPDEA, that our lab previously detected as a cytosolic PDE in wild type epimastigotes. Supplemental figure 1) or TcrPDEC2 phosphodiesterase (a phosphodiesterase associated to the contractile vacuole complex and involved in osmotic stress response (Schoijet et al., 2011)) genes, to produce constitutive overexpressing cell lines for these enzymes. After the selection period, we confirmed the overexpression of TcrPDEC2 by evaluating the PDE activity associated to membranes and TcrPDEA1 expression by western blot (Figure 3, C and D). Under standard culture conditions, transfected parasites showed no significant difference in growth compared to a wild type culture. We then proceeded to test the proliferation capacity of these transfected parasites after oxidative stress, in comparison to a control line containing only the empty vector. Figure 3B shows that parasites overexpressing TcrPDEC2 could proliferate at the same rate as control parasites after exposure to different peroxide concentration. However, TcrPDEA1 overexpressing parasites showed a statistically significant decrease in Thymidine incorporation when exposed to a sub-lethal concentration of hydrogen peroxide, accounting for a diminished replication.

**Figure 3.**
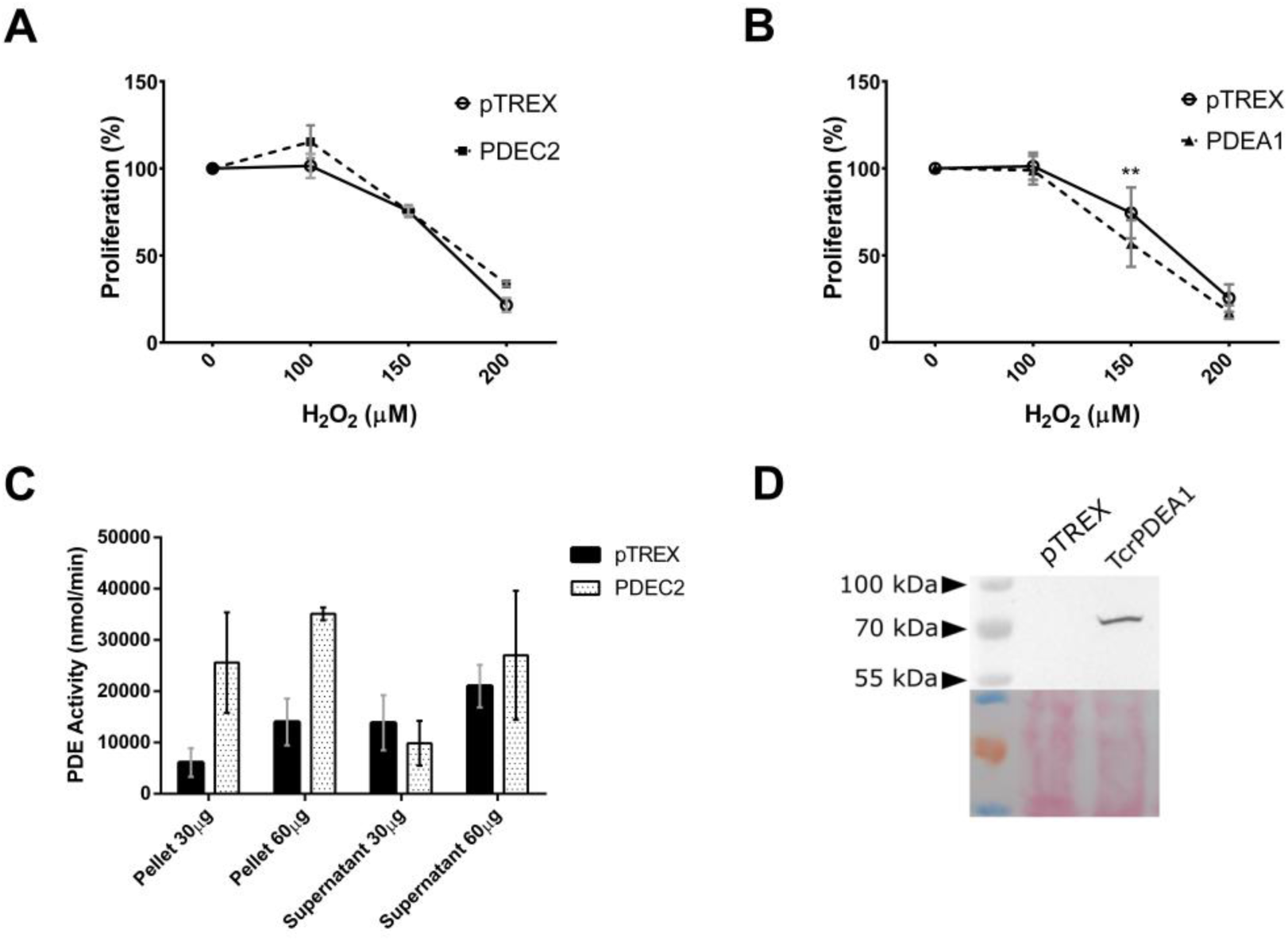
TcrPDEA1 overexpression causes reduced proliferation under oxidative stress. **(A)** Overexpression of TcrPDEC2 has no effect on proliferation under stress (n = 3). **(B)** In contrast, overexpression of TcrPDEA1 causes reduced resistance and proliferation under H_2_O_2_ treatment (n = 3). Statistical analysis is a Two-way ANOVA with Sidak’s multiple comparison test. Error bars represent Standard Error of the Mean. **(C)** Overexpression of the phosphodiesterases was confirmed by PDE activity assays (representative assay, n = 3, error bars represent Standard Deviation) or **(D)** Western blot with a polyclonal antibody against TcrPDEA1 (representative assay, n = 2). Bottom panel shows the Ponceau staining as loading control. (**) p<0.01

### Cyclic AMP analog improve infection of trypomastigotes in Vero cells

Trypomastigotes also face the challenge of coping with oxidative stress, delivered in this instance by the early innate immune response of the mammalian host. The initial oxidative burst releases ROS and RNS that aim to eliminate the invading parasites, but also function as external signal for the parasites to modulate the infection (Paiva et al., 2018).

According to the above results, epimastigotes showed a higher viability under oxidative stress conditions when pre-treated with cAMP analogs. We aimed to test if this second messenger was also involved in this response in the infective stage (trypomastigote). To this end, we used trypomastigotes of the Tulahuen strain that express the recombinant *E. coli* β-galactosidase. This allows the estimation of the number of amastigotes in an infected culture and thus a quantification of the magnitude of the infection by measuring the increase in the absorbance of the lysate. Trypomastigotes were treated with 8-pCPT-cAMP for 30 min, while being exposed to hydrogen peroxide simultaneously during the last 10 min of incubation, as described in Materials and Methods (see also schematic diagram in Figure 4A). Infection was allowed to proceed for 48 h and infection levels evaluated as described under Materials and Methods.

**Figure 4.**
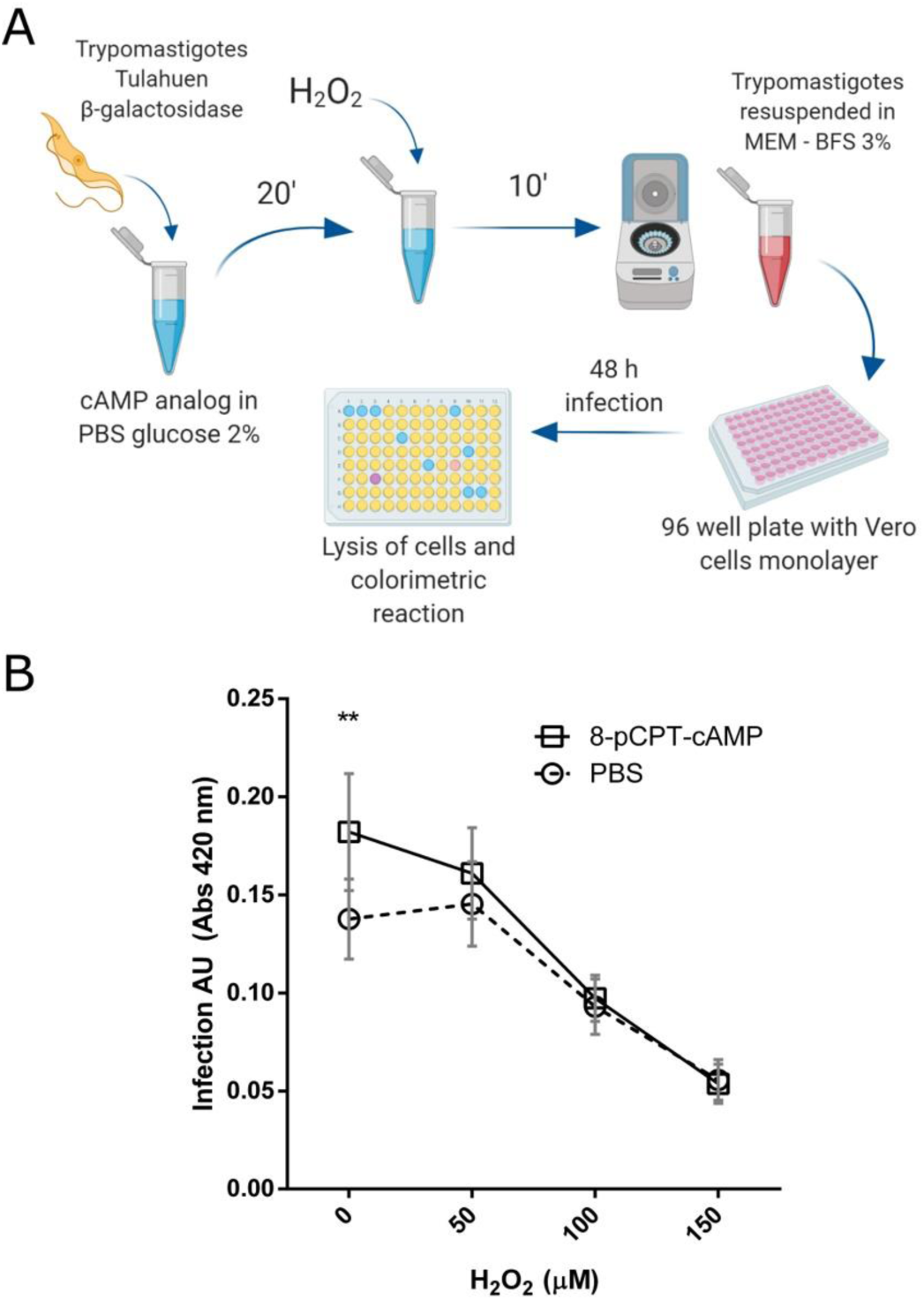
Pre-incubation with cAMP can enhance infection of trypomastigotes *in vitro*. **(A)** Schematic representation of the experimental protocol as described in Materials and Methods. Trypomastigotes where exposed to cAMP analog for 30’ or/and hydrogen peroxide for the final 10’ before infection of Vero cells. Treated trypomastigotes were removed from culture after 24 h and infection was allowed to develop for up to 48 h. **(B)** Infection measured as Beta-galactosidase colorimetric reaction. Values represent the mean of nine independent experiments. Error bars represent standard error of the mean. A Two-way ANOVA analysis established that the analog has a significant effect over amastigote production (p value = 0.0143).

Figure 4B shows that when trypomastigotes are treated with increasing concentrations of hydrogen peroxide before infection, fewer amastigotes are detected in a dose dependent-manner. It has been reported that mild oxidative stress, as with other types of stress, can enhance viability or infectivity in trypanosomatids (de Almeida Nogueira et al., 2011; Goes et al., 2016; Nogueira et al., 2015; Paes et al., 2011). Our results show this could also be the case for doses lower than 50 μM H_2_O_2_. However, cAMP increase, before stress exposure, seems to alter this effect, neutralizing the slight increase in infectivity. Still, cAMP analog treatment alone, increased amastigotes production in a significant manner.

## Discussion

The cAMP pathway and related proteins have been proven to participate in several cellular processes of the trypanosomatid parasites. Examples of this are the “quorum sensing” like behavior of the *Trypanosoma brucei* culture (Lopez et al., 2015), the lethal effect of the RNAi of the phosphodiesterases TbPDEB1 and TbPDEB2 in this same organism (Oberholzer et al., 2007), and the differentially expressed LdPDEA role in the oxidative stress response of *Leishmania donovani* (Bhattacharya et al., 2009), amongst others. In *T. cruzi*, TcrPDEC2 has been involved in the osmotic stress response named “Regulatory Volume Decrease” (Schoijet et al., 2011). Also, intracellular cAMP concentration increases under differentiation conditions while cAMP analogs and PDEs inhibitors could induce differentiation (Fraidenraich et al., 1993; Gonzales-Perdomo et al., 1988; Rangel-Aldao et al., 1988). All this evidence indicates that the cAMP transduction pathway is regulating numerous cellular responses indispensable for the correct progression of the life cycle of the trypanosomatids. In this work, we showed that, cAMP intracellular levels regulate *T. cruzi* resistance to oxidative stress. Different cAMP analogs improved epimastigotes proliferation after suffering oxidative stress with sub-lethal concentrations of hydrogen peroxide, while these same analogs had no significant effect under control conditions. The consistency of this effect depends on the lipophilicity, stability and specificity for the PKA and/or for other effectors of the cAMP analogs, which was also observed in similar assays performed on *Leishmania donovani* (Bhattacharya et al., 2009). Moreover, the overexpression of the cytoplasmic phosphodiesterase TcrPDEA1, an ortholog of the LdPDEA, turned the parasites more vulnerable to these same peroxide concentrations. The fact that overexpression of cytosolic TcrPDEA1 affected resistance but TcrPDEC2, localized in the contractile vacuole, didn’t, supports the theory that the effect of cAMP is regulated in the micro-domains where the adenylyl cyclases, phosphodiesterases and cAMP effector proteins co-localize, and for that, a generalized increase in intracellular cAMP wouldn’t trigger all the transduction pathways where it can influence.

The experiments performed on trypomastigotes and Vero infections in this work leave an open possibility that oxidative stress response and infectivity are regulated by crosstalking transduction pathways. Our results do not allow us to reject the hypothesis that cAMP could improve viability of the trypomastigotes during the infection process, or infectivity, or both, in the absence of oxidative stress. But, since parasites treated with a cAMP analog respond in a non-synergic different way to sub-lethal doses of hydrogen peroxide, compared to the vehicle treated parasites, these transduction pathways might possibly share a common node, or several. Nevertheless, we conclude that in this stage, cAMP doesn’t increase trypomastigote resistance to oxidative stress.

These observations could be explained by a model similar to the one Bhattacharya et al proposed for *Leishmania*, in which during the vector early stages of the parasite life cycle the cAMP transduction pathway is aimed to deal with the oxidative environment and cAMP levels are maintained under a certain threshold by the cytosolic PDEA. But as the redox status changes closer to the urinary tract of the vector, the cAMP is accumulated in higher quantities due to reduction of PDEA expression, and triggers different mechanisms, now associated with infectivity and antioxidant molecules accumulation to deal with the oxidative burst of the immune system in the host. It would be interesting to investigate the TcrPDEA1 behavior during differentiation to define whether this model is replicated in *T. cruzi* life cycle.

Redox status has recently been found to act as another possible environmental signal for the trypanosomatid parasites to define which metabolic mechanisms to put in action through its life cycle. Nogueira et. al. presented results that show how modulation of the redox status in the environment, even inside the vector’s midgut, can change the fate of the epimastigotes of *T. cruzi*, inducing proliferation or differentiation (Nogueira et al., 2015). These findings underscore the importance of understanding the mechanisms by which this parasite senses the redox status and responds to cope with the environmental stress ahead.

In summary, our results demonstrate for the first time that cAMP is involved in the oxidative stress response in the epimastigote stage of *T. cruzi*, and we propose TcrPDEA1 as the main phosphodiesterase regulating this process and cAMP transduction pathway as a participant of redox status sensing in *T. cruzi*. Further experiments should be performed to unravel how TcrPDEA1 modulates this response and which other proteins are involved. Also, our results confirm the previous observations that cAMP can improve infection capability. Additionally, we show for the first time the crosstalk between the cAMP transduction pathway and oxidative stress response to hydrogen peroxide in trypomastigotes *in vitro*.

## Acknowledgments

This work was supported by the Consejo Nacional de Investigaciones Científicas y Técnicas (PIP 2013-00351); Departamento de Fisiología, Biología Molecular y Celular, Facultad de Ciencias Exactas y Naturales, Universidad de Buenos Aires (UBACyT 2014-2017, Nro 00155BA); and Agencia Nacional de Promoción Científica y Tecnológica (PICT 2013–2015, PICT 2015-0898 and PICT 2017-2125). G.D.A. and A.C.S. are members of the Research Career of Consejo Nacional de Investigaciones Científicas y Técnicas, T.S. is a fellow from the same institution. The funders had no role in study design, data collection and analysis, decision to publish, or preparation of the manuscript. The authors have no conflict of interest to declare.

