## Supplemental Figure 1 for "Intracellular cyclic AMP levels modulate differential adaptive responses on epimastigotes and cell culture trypomastigotes of *Trypanosoma cruzi*"

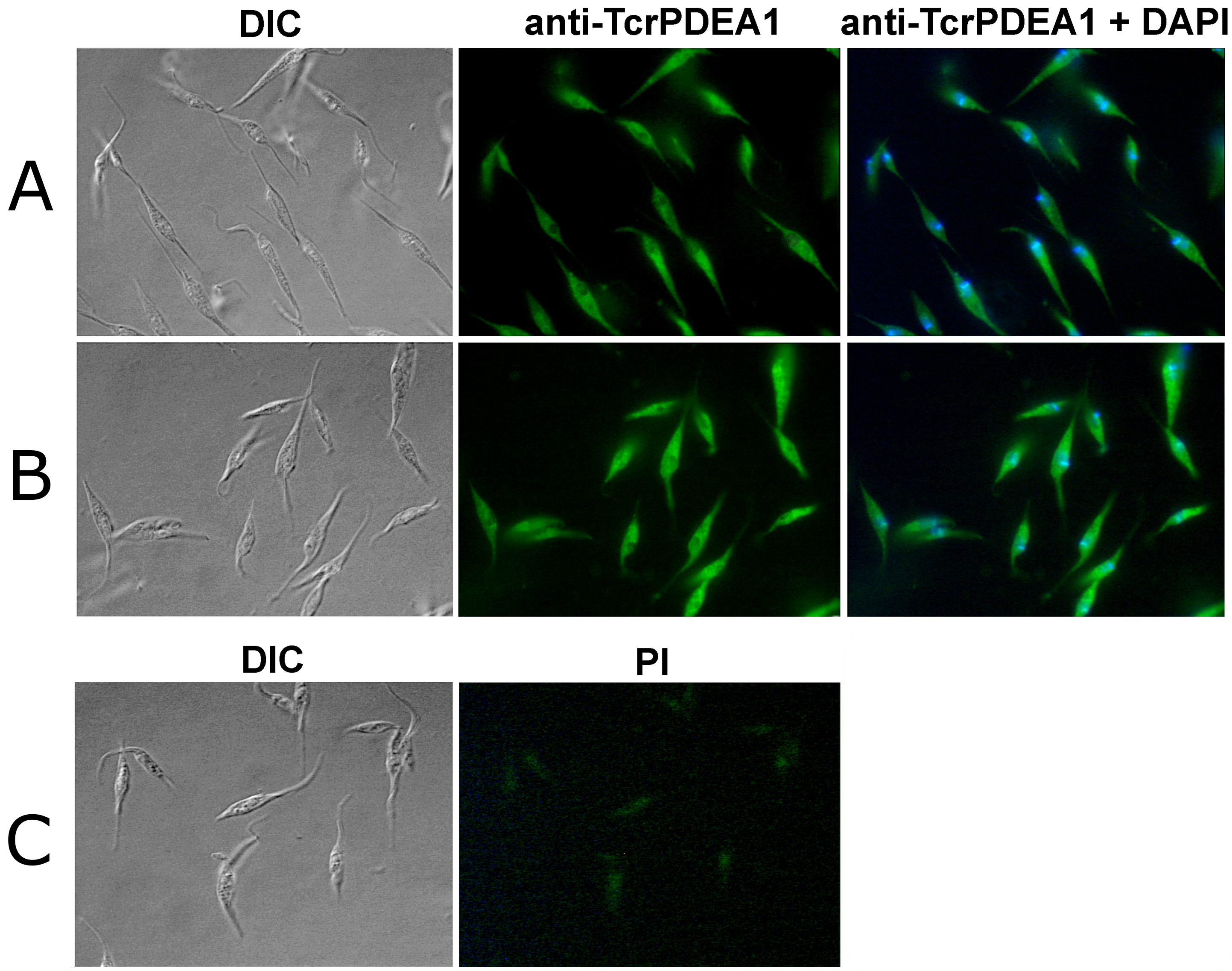


Figure S1. **TcrPDEA1 is a cytosolic phosphodiesterase.** (A) and (B) show an indirect inmmunofluoresce of CL brener wild-type epimastigotes fixed and stained with anti-TcrPDEA1 antibody (green) and DNA staining with DAPI (blue). (C) shows background fluorescence of the negative control (secondary antibody only). DIC corresponds to differential interference contrast.
